# Multiplexed detection of RNA using MERFISH and branched DNA amplification

**DOI:** 10.1101/505784

**Authors:** Chenglong Xia, Hazen P. Babcock, Jeffrey R. Moffitt, Xiaowei Zhuang

**Affiliations:** Howard Hughes Medical Institute, Department of Chemistry and Chemical Biology, and Department of Physics, Harvard University, Cambridge, MA 02138, USA; Center for Advanced Imaging, Harvard University, Cambridge, MA 02138, USA; Program in Cellular and Molecular Medicine, Boston Children’s Hospital; Department of Microbiology, Harvard Medical School, Boston, MA 02115, USA

## Abstract

Multiplexed error-robust fluorescence in situ hybridization (MERFISH) allows simultaneous imaging of numerous RNA species in their native cellular environment and hence spatially resolved single-cell transcriptomic measurements. However, the relatively modest brightness of signals from single RNA molecules can become limiting in a number of important applications, such as increasing the imaging throughput, imaging shorter RNAs, and imaging samples with high degrees of background, such as some tissue samples. Here, we introduce a branched DNA (bDNA) amplification approach for MERFISH measurements. This approach produces a drastic signal increase in RNA FISH samples without increasing the fluorescent spot size for individual RNAs or increasing the variation in brightness from spot to spot. Using this approach in combination with MERFISH, we demonstrated RNA imaging and profiling with a near 100% detection efficiency. We further demonstrated that signal amplification improves MERFISH performance when fewer FISH probes are used for each RNA species, which should allow shorter RNAs to be imaged. We anticipate that the combination of bDNA amplification with MERFISH should facilitate many other applications and extend the range of biological questions that can be addressed by this technique in both cell culture and tissues.

## Introduction

Single-molecule fluorescence in situ hybridization (smFISH) provides both quantitative measurements of RNA expression and information about RNA spatial localization by directly imaging individual RNA molecules in single cells^1, 2^. The ability of smFISH to visualize gene expression at single-cell resolution has generated many critical insights for different biological processes, such as cell fate determination during cell division, local translation, cell migration, the establishment of cell polarity, and body patterning in development^3^. In recent years, multiplexed smFISH^4–9^ and in situ sequencing^10–12^ have been developed to increase the number of RNA species that can be simultaneously imaged within cells or tissues, with several technologies enabling the profiling of hundreds to thousands of RNAs simultaneously at single cell resolution^9, 11–13^. These approaches have been used to reveal the internal organization of the transcriptome within cells, discover novel cell types and identify cells based on their expression profile, and map out the organization of different cell types within tissues^9, 12–14^. Among these approaches, multiplexed error robust fluorescence in situ hybridization (MERFISH) massively multiplexes smFISH by assigning error-robust barcodes to individual RNA species, labeling RNAs with oligonucleotides that represent each barcode, and sequential smFISH imaging to read out these barcodes^9^, which has allowed single-cell transcriptomic profiling in both cultured cells or tissue slices^9, 14, 15^.

The fluorescent signal produced from a single RNA molecule in MERFISH is generated from the binding of many fluorescently labeled probes to each RNA. While the signal produced from these probes is sufficiently bright to allow individual RNA molecules to be identified and detected in cell culture^9, 16^ and cleared slices of the mouse brain^14, 15^, the relatively modest brightness of these signals makes some biological questions still challenging to address. For example, limited signal brightness requires relatively long camera exposures with high-power laser illumination sources, which in turn limits the number of cells and the volume of tissue that can be imaged in a given time. Moreover, the use of multiple unique probes per RNA requires targeted RNAs to have sufficient length to bind these probes, limiting the measurement of shorter RNAs. Finally, although a customized clearing approach has been reported for MERFISH^15^, residual autofluorescence, light scattering, and autofluorescent granules that are difficult to remove (e.g. lipofuscin^17^) can produce signals that challenge the imaging of single RNA molecules in some samples.

Each of these limitations could be addressed by using amplification to increase the brightness of the signal produced via individual RNA molecules; thus, it is desirable to combine MERFISH with methods for signal amplification. In addition to the degree of signal increase, there are other important properties that should be considered for the ideal amplification method. First, to accurately detect and distinguish closely spaced RNAs, it is desirable to have an amplification method that does not substantially increase the fluorescent spot size after amplification. Next, to more easily differentiate background spots from spots produced by real RNAs, it is desirable not to substantially increase the spot-to-spot brightness variation after amplification. Last, MERFISH utilizes multiple distinct readout sequences, one associated with each bit in the barcodes assigned to RNAs; thus, the amplification method should be easily extended to the amplification of a large number of orthogonal sequences.

Several methodologies have already been introduced to amplify FISH signals. In hybridization chain reaction (HCR), two fluorescently labeled metastable hairpin oligonucleotides self-assemble into long fluorescent polymers starting from an initiator sequence present on each probe^18–20^. In rolling-circle amplification (RCA), each probe is circularized to create a template which is then replicated into long concatenated copies containing binding sites for a second fluorescently labeled oligonucleotide probe^21–23^. Finally, in branched DNA (bDNA) amplification, primary and secondary amplifier oligonucleotides, each containing multiple replicate binding sites, are assembled on each smFISH probe to form a branched structure which binds multiple copies of a fluorescently labeled probe^24–27^. Recently, a bDNA-like approach was introduced, clampFISH, in which the amplifier molecules in each round of amplification are covalently circularized via click chemistry, topologically entangling individual amplifier molecules to increase binding specificity^28^.

Each technique has advantages and potential disadvantages. HCR and RCA have the advantage that the degree of amplification can be tuned by changing the hybridization or polymerization time and a very large degree of amplification can be obtained. However, both techniques have been reported to increase the size of fluorescent spots, from the diffraction limit up to ∼1 micron^20, 29^. In addition, HCR has been reported to introduce a variable degree of amplification for different copies of the same target molecule (with a variation of ∼10-fold)^13^. By contrast, the degree of amplification in bDNA amplification is fixed by design, i.e. the assembled bDNA structures cannot grow indefinitely, even in the presence of abundant reagents. We thus anticipate that it will be more straightforward to control the spot size or limit the increase in variability in brightness from molecule to molecule with this approach. However, only a small number of primary and secondary amplifier pairs have been reported, and bDNA has not yet been combined with highly multiplexed RNA imaging.

Here, we developed an approach that combines MERFISH with bDNA amplification to dramatically increase the MERFISH signal brightness. We demonstrated that bDNA amplifiers can be rapidly assembled, that they can increase the spot brightness by ∼30-fold without increasing spot size, and that they do not increase the variability in spot brightness. To combine this approach with MERFISH, we developed 16 pairs of orthogonal bDNA amplifiers and then demonstrated with these amplifiers substantial increase in single-molecule signals in MERFISH measurements. Although the MERFISH performance was similar with and without amplification when a large number of hybridization probes were used per gene, when fewer hybridization probes were used, bDNA amplification substantially improved the detection efficiency of MERFISH. This improvement should allow MERFISH to be extended to the study of short RNAs. bDNA amplification thus promotes high performance MERFISH measurements by providing a fast, simple, and efficient way to simultaneously amplify the signal from a large number of RNAs in single cells.

## Results

### The design of a bDNA amplification scheme for MERFISH measurements

In MERFISH, each RNA of interest is assigned a unique barcode drawn from a barcoding scheme that enables error detection and, when needed, error correction. The sample is then stained with a complex library of oligonucleotide probes, termed encoding probes, which effectively label each RNA species with the desired barcode^9^,^16^. Each encoding probe contains a 30-mer ‘target region’, whose sequence is complimentary to a region of the target RNA, and multiple 20-mer ‘readout’ sequences (Fig. 1a). For binary barcodes, there is one unique readout sequence per bit in the barcode, and the encoding probe set corresponding to a given RNA contains the readout sequence for a given bit if the barcode assigned to that RNA contains a ‘1’ in that bit. The presence or absence of a ‘1’ in a given bit is then determined by hybridizing a fluorescently labeled ‘readout probe’ complementary to the corresponding readout sequence.

**Figure 1.**
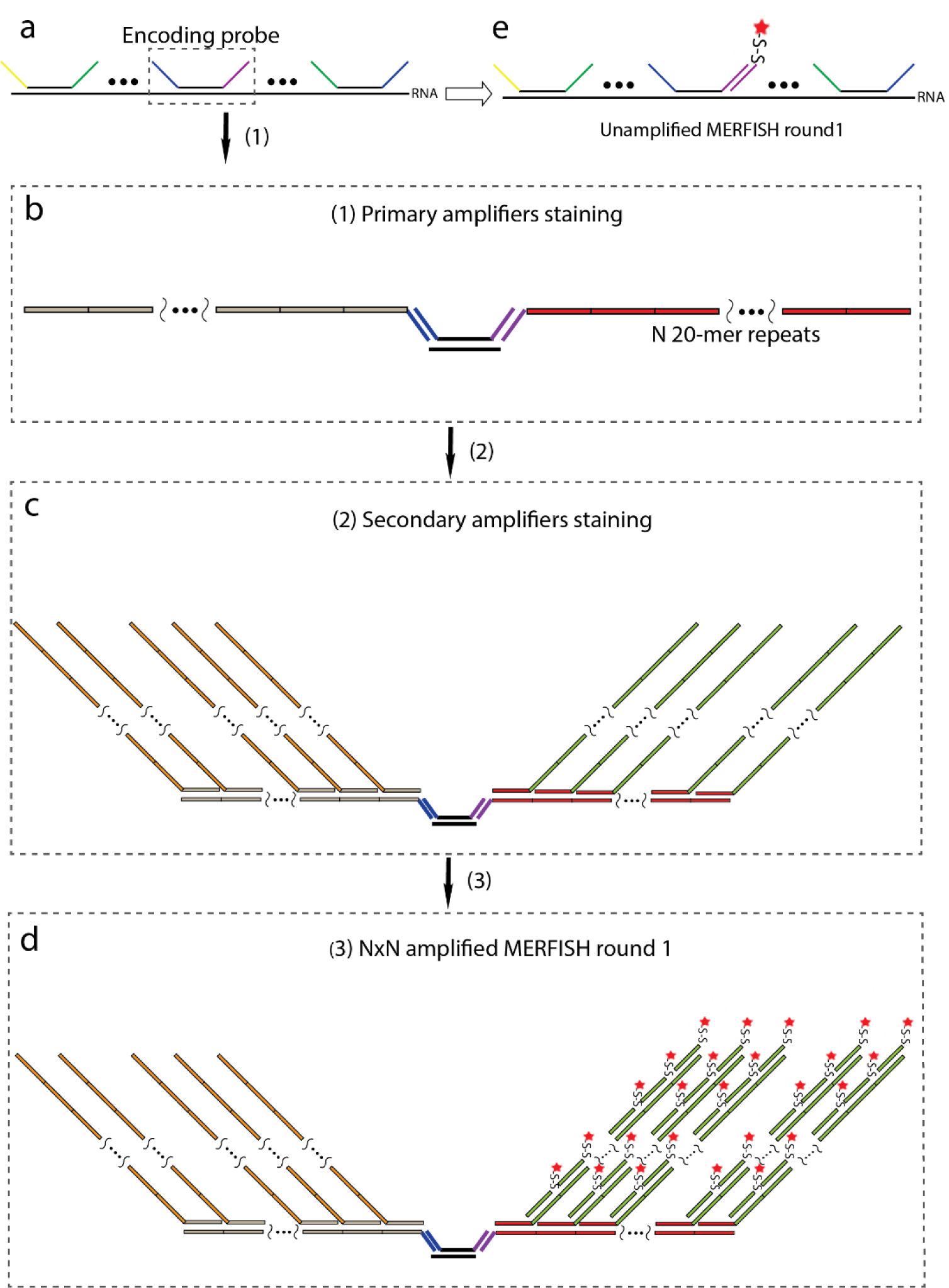
Schematic illustration of bDNA amplification for MERFISH imaging. (a) Depiction of a tile of multiple MERFISH encoding probes bound to a single RNA molecule. Each encoding probe has a 30-mer target sequence (black line) and multiple readout sequences (yellow, green, purple, and blue lines represent different readout sequences). (b) Schematic depiction of the binding of two primary amplifier oligonucleotides to their corresponding readout sequences for the encoding probe in the dashed box in (a). The primary amplifiers have a complimentary sequence to the readout sequence on encoding probes (blue or purple lines) and N 20-mer repeating sequences unique to each primary amplifier (tan or red lines). (c) Schematic depiction of the binding of secondary amplifiers to the primary amplifier’s repeating regions. The secondary amplifiers have a complimentary sequence to one of the 20-mer repeating sequences on the primary amplifiers (tan or red lines) and N 20-mer repeating sequences that are complimentary to a unique readout probe for each secondary amplifier (orange or green lines). (d) Schematic depiction of the binding of a fluorescently labeled readout probe in a N×N amplified specimen after the first round of MERFISH readout. There is a one-to-one correspondence between the original readout sequence on the encoding probe and the final readout sequence repeated on the bDNA structure bound to that readout sequence. (e) Schematic depiction of the first round of MERFISH readout staining without amplification.

Our strategy to incorporate bDNA amplification with MERFISH is summarized in Figure 1. First, we bind to the sample MERFISH encoding probes as described above. We then bind a set of primary amplifier oligonucleotides, in which a unique primary amplifier is targeted to each readout sequence (Fig. 1b). Each primary amplifier contains *N* repeats of a unique 20-mer binding site. A set of secondary amplifier oligonucleotides are then added to the sample (Fig. 1c). Each unique secondary amplifier is targeted to the binding site on one corresponding primary amplifier and contains *N* repeats of another unique 20-mer binding site (Fig. 1c). To readout the bit associated with a given readout sequence, we then hybridize to the sample a fluorescently labeled probe targeting the binding site on the secondary amplifier associated with that readout sequence (Fig. 1d). In this fashion, each original readout sequence would correspond to N^2^ copies of another binding site unique to that readout sequence. Thus, the effective readout signal would be theoretically amplified N^2^ fold. We term this process N×N amplification.

In addition, we made one notable modification to our design of the bDNA amplifiers as compared to those previously reported^24, 30^ to potentially improve the performance of these amplifiers. Specifically, we designed the amplifiers sequences using only three of the four nucleotides. Because probe sequences that contained only three of the four nucleotides have been previously shown to contain significantly less secondary structures than sequences that used all four nucleotides^31^, we anticipate that these amplifiers will have higher hybridization rates and high probability of assembling correctly and efficiently. Indeed, we have previously shown that readout probes designed using a reduced nucleotide alphabet have these properties^16^.

### Properties of bDNA amplification

To test this strategy and quantify the performance of bDNA amplification on readout sequences, we first designed one pair of three-letter bDNA amplifiers and performed amplification on smFISH probes targeting a single RNA. The smFISH probes were designed in a similar way to MERFISH encoding probes: each smFISH probe contains a 30-mer target sequence complimentary to the target RNA template and four different 20-mer readout sequences used in MERFISH measurements. We only used one of these readout sequences for this smFISH measurement. We designed 48 encoding probes targeting different regions of the filamin A (FLNA) mRNA. We then stained a culture of human osteosarcoma cells (U-2 OS) cells with these probes. To reduce background, we utilized a matrix imprinting and clearing approach in which these samples were also stained with an acrydite-modified poly(dT) locked nucleic acid (LNA) oligonucleotide ‘anchor probe’ that targets the poly(A) tail of mRNAs^15^. We then embedded these samples in a thin film of polyacrylamide to which the anchor probes were covalently incorporated, then utilized detergents and proteinase K to clear the sample of lipids and proteins.

Following the clearing, we then stained these samples with the primary amplifier for 15 minutes. To preserve the binding of encoding probes during amplifier staining and washing steps, we used hybridization conditions suitable for the 20-mer amplifier binding sites but in which the 30-mer target regions would remain stably bound: 10% formamide at 37 °C. We then washed the sample for 15 minutes in the same conditions to remove excess primary amplifier and repeated this process with the secondary amplifier. Each amplifier contained 5 binding sites in this 5×5 amplification scheme. We then measured the average FLNA smFISH spot brightness of samples either directly labeled with readout probes on the encoding probes (unamplified) or labeled with readout probes targeting the amplified bDNA (amplified) (Fig. 2a,b). We observed a 10.5-fold signal increase averaged across ∼10,000 molecules in the amplified scheme versus the unamplified detection scheme (Fig. 2c). This increase represents 42% of the theoretical maximum amplification value.

**Figure 2.**
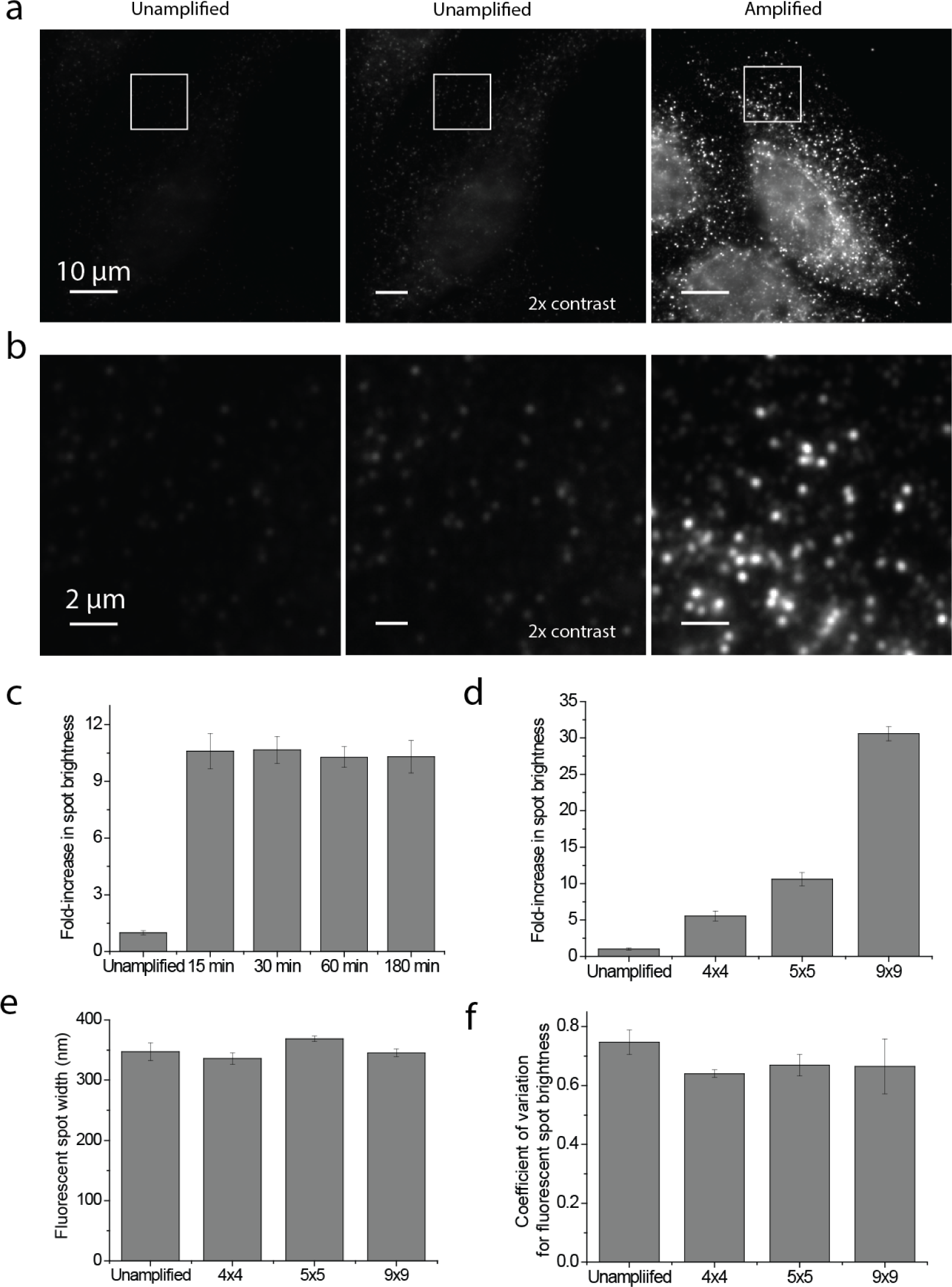
bDNA amplification dramatically increases signal brightness without increasing spot size or variation in brightness from spot to spot. (a) Images of U-2 OS cells stained with smFISH probes to the filamin A mRNA (FLNA). Left: an unamplified sample stained with a Cy5-labeled readout probe. Middle: the same image as on the left but with a 2x increased contrast to better illustrate the fluorescence signal. Right: a sample 5×5 amplified and stained with a Cy5-labeled readout probe. Scale bars, 10 µm. (b) Zoom in of white boxed region in (a). Scale bars, 2 µm. (c) The fold-increase in average brightness of individual FLNA mRNA spots after staining with amplifiers for different durations relative to the brightness observed for unamplified samples. (d) The fold-increase in average brightness of individual FLNA mRNA spots in 4×4 amplified, 5×5 amplified, and 9×9 amplified samples relative to the brightness observed for unamplified samples. The value reported for 5×5 amplification was reproduced from the 15 min measurement in (c). (e) The average FLNA mRNA spot sizes in unamplified, 4×4 amplified, 5×5 amplified, and 9×9 amplified cells. The width (full width at half maximum) was determined by Gaussian fitting of RNA spots. (f) The coefficient of variations for RNA spot brightness in unamplified, 4×4 amplified, 5×5 amplified and 9×9 amplified samples. Error bars in (c) – (f) represent standard deviation across three replicates.

We reasoned that one possibility we did not observe the theoretical maximum amplification factor was that the amplifier staining time was not sufficient and that the bDNA constructs had not fully assembled. To test this possibility, we conducted a time series of the same 5×5 amplification, with increasing amplifier staining time. We found that the amplified signal was already saturated when we hybridized amplifiers with 15 minutes each round (Fig. 2c).

Next, we asked whether the number of repeating sequences on each amplifier could change the amplification performance. We designed a pair of 4×4 amplifiers and 9×9 amplifiers using the same binding sequences in the 5×5 amplifiers described above and repeated the amplification with FLNA smFISH probes (Fig. 2d). We observed a 5.5-fold and a 30.6-fold signal increase with 4×4 amplification and 9×9 amplification, respectively. Thus, the degree of amplification can be tuned with the number of binding sites, and for all amplification schemes tested we observed ∼40% of the theoretical amplification values.

One potential consequence of amplification is the increase of FISH spot sizes, which could limit the density of RNAs that can be imaged and identified due to an increased chance of the signal from one molecule overlapping with that from another molecule. As a rough estimate of the potential spot size increase due to bDNA amplification, we considered the length of a fully extended 9×9 amplifier scaffold which is expected to be 132 nm. This length is within the diffraction limit; thus, we did not anticipate a measurable increase in the spot size with even the largest amplification considered. Indeed, the measured spot sizes of unamplified, 4×4 amplified, 5×5 amplified, and 9×9 amplified samples were identical (Fig. 2e).

Another concern for amplification approaches is the potential to increase the variability in brightness from one molecule to another based on differential amplification. In principle, the finite amplification provided by the defined assembly of the bDNA structures should limit this variability, as the assembly reaction can be run to completion or saturation. To determine any potential increase in the variation in spot brightness due to amplification, we measured the coefficient of variation in spot brightness for unamplified, 4×4 amplified, 5×5 amplified, and 9×9 amplified samples. Notably, we found that the coefficient of variation is similar for all degrees of bDNA amplification, indicating that this approach does not increase the variation in spot brightness beyond that observed for unamplified samples (Fig. 2f).

### Amplifier screening for MERFISH imaging

To extend bDNA amplification to MERFISH measurements, it is necessary to have a unique primary and secondary amplifier pair for each readout sequence used in the measurement. For example, with a previously published 16-bit, modified Hamming distance-4 (MHD4) encoding scheme^9^, 16 pairs of primary amplifiers and secondary amplifiers are needed. However, previous applications of bDNA have only reported a few amplifier pairs^24, 26, 30^ and the reported pairs did not utilize the three letter alphabet; thus, it was necessary for us to design a large set of new amplifier pairs. To this end, we anticipated that the lower probability of unanticipated secondary structure provided by the use of three-letter sequences would be beneficial.

We designed random 20-mer, three-letter repeating sequences with a per-base probability of 25% for A, 25% for T, and 50% for C or 25% for A, 25% for T, and 50% for G for primary amplifiers and secondary amplifiers, respectively. We selected from these sequences a set of orthogonal sequences with limited cross homology using a previously described algorithm^32^ and then blasted these sequences to the human transcriptome to avoid homology regions longer than 11 nucleotides, as described previously^16^. We designed a set of orthogonal 5×5 amplifier pairs with 20-mer, three-letter sequences and screened each of these pairs for its ability to amplify smFISH signals. We found that 80% of the amplifier pairs designed worked as predicted, producing uniform, bright signals. The remaining 20% of the amplifiers revealed two types of defects (Fig. 3). First, we observed that a small number of secondary amplifiers bound to other cellular components than our probes (Fig. 3a, b). These signals were RNase-dependent indicating that these amplifiers were binding non-specifically to cellular RNA (Fig. 3c). Given the cellular distribution of this binding, we suspected these guanine-rich secondary amplifier sequences might form G-quadruplex structures with mitochondrial RNAs^33^. In parallel, we observed a small number of amplifier pairs that assembled with much lower efficiency, producing only small degrees of amplification, perhaps due to low melting temperature or the formation of G-quadruplex structures that inhibit proper amplifier assembly (Fig. 3e). When we replaced these amplifiers with new pairs containing different sequences, both the high background problem (Fig. 3d) and the low amplification efficiency problem were solved (Fig. 3f). Thus, from 20 pairs of 5×5 amplifiers, we identified 16 suitable for MERFISH imaging.

**Figure 3.**
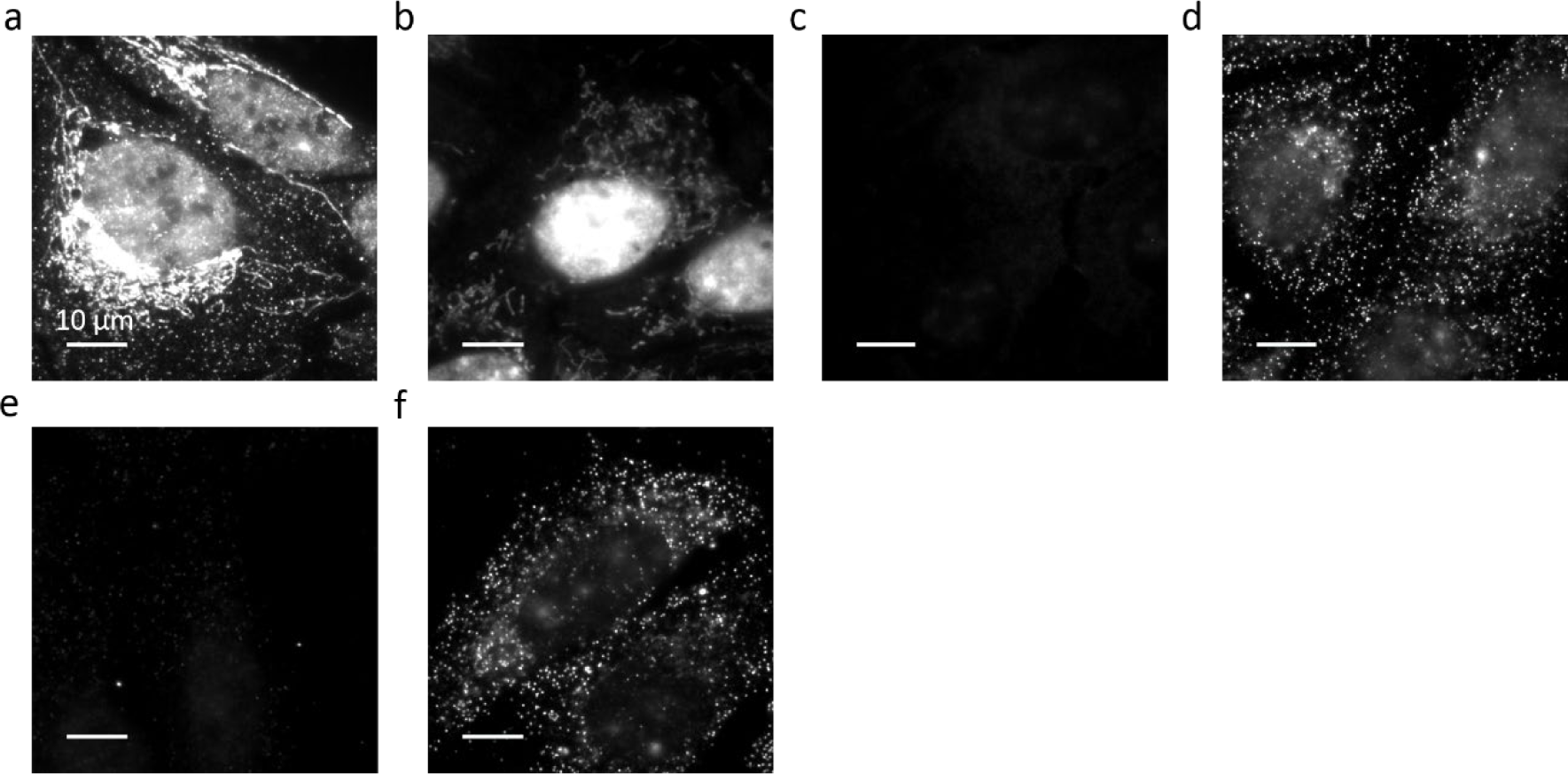
Examples of amplifiers with high background or low amplification efficiency during amplifier screening. (a) A U-2 OS sample stained with an amplifier pair that produced high background. (b) As in (a) but using only the secondary amplifier used in (a). (c) As in (a) but for a sample treated with RNase prior to staining with amplifiers. (d) A U-2 OS sample stained with a different amplifier pair that targets the same readout as the pair used in (a) and which does not show the high background of (a). (e) A U-2 OS sample stained with a pair of amplifiers showing low amplification efficiency (1.25-fold increase). (f) A U-2 OS sample stained with a different amplifier pair that targets the same readout as the pair used in (e) but with substantially high amplification than observed in (e). All images are displayed with the same contrast. Scale bars, 10 µm.

### MERFISH measurements with bDNA amplification

To determine if the 16 pair of amplifiers we identified work with MERFISH measurements, we performed 5×5 amplification on a 130-RNA library that has been previously measuring using MERFISH without amplification and which showed both high accuracy and high detection efficiency^16^. We performed MERFISH imaging in U-2 OS cells, and used 8 rounds of two-color imaging to read out 16 bits, as well as reductive cleavage of disulfide bonds to remove the fluorophores linked to the readout probes between consecutive rounds of imaging for both amplified and unamplified samples, as described previously^16^. Figure 4a-c shows that individual RNA molecules could be clearly detected in each of the eight hybridization and imaging rounds of 5×5 amplified samples, allowing their identity to be decoded.

**Figure 4.**
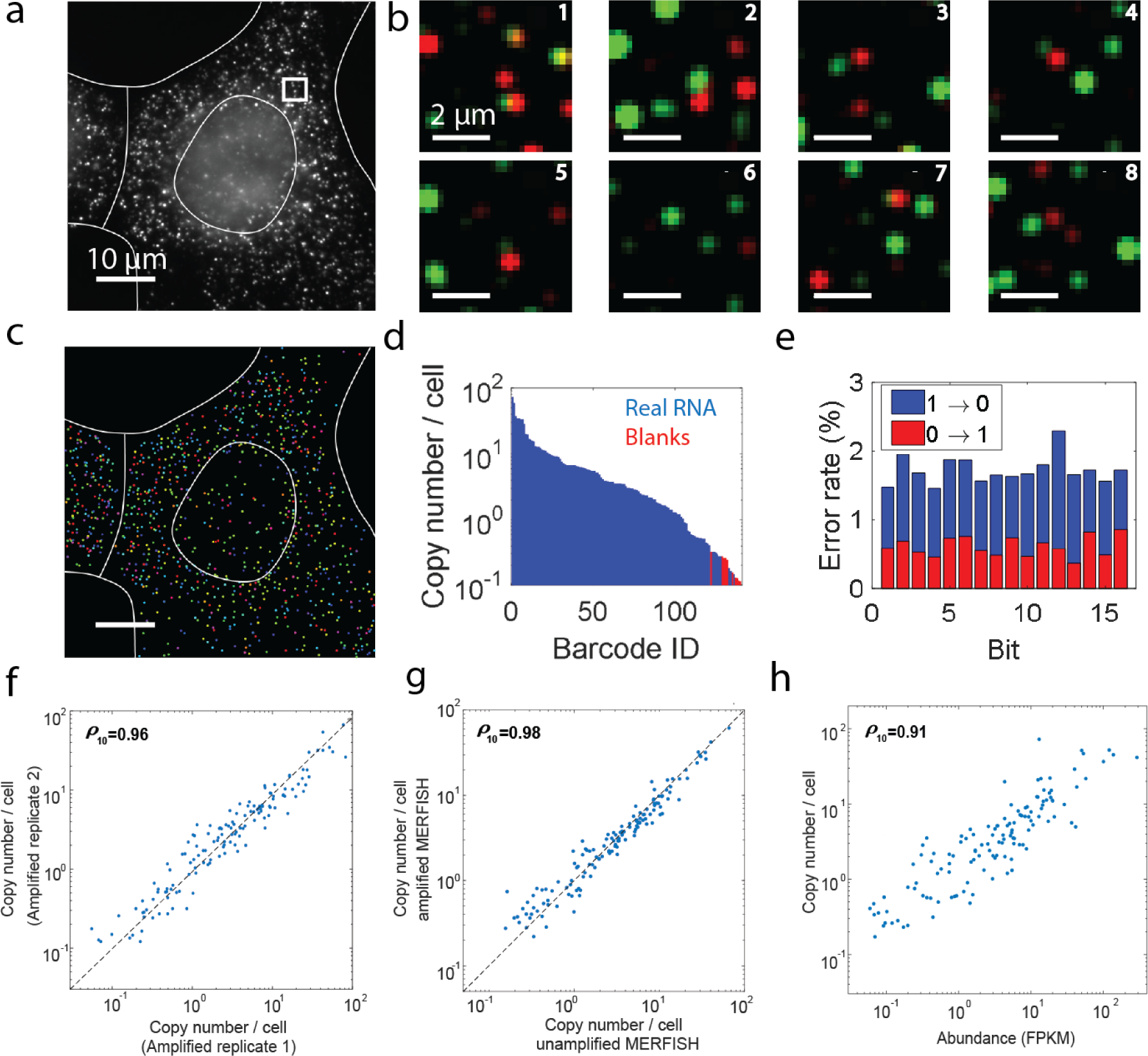
MERFISH measurements of 130 RNAs with 5×5 amplification in U-2 OS cells. (a) Image of a 5×5 amplified sample stained with encoding probes for 130 RNA species and a Cy5-labeled readout probe that detects one of the bits of the RNA barcodes. Scale bar, 10 µm. (b) Two-color smFISH images from each of the eight rounds of hybridization and imaging using readout probes labeled with Cy5 (green) or Alexa750 (red) for the white boxed region in (a). Scale bars, 2 µm. (c) All identified RNAs detected in the region depicted in (a) with the barcodes of the RNAs represented by the colors of the markers. Scale bar, 10 µm. (d) The average RNA copy numbers per cell for real RNA barcodes (blue) and the blank control barcodes (red) detected with 5×5 amplification, sorted from largest to smallest value. (e) The error rates, the fraction of measured barcodes that contain a given bit flip, for each bit with 5×5 amplified MERFISH measurements. The 1-to-0 error rates are depicted in blue and the 0-to-1 error rates are depicted in red. (f) The average copy numbers per cell detected in one 5×5 amplified sample versus a replicate sample. The Pearson correlation coefficient is 0.96. (g) The average copy numbers per cell observed for each RNA species in 5×5 amplified U-2 OS cells versus the copy numbers obtained without amplification. The Pearson correlation coefficient is 0.98. The copy number per cell values were averaged across three replicates each of amplified and unamplified samples. (h) The average RNA copy numbers per cell determined by MERFISH with 5×5 amplification versus the abundances determined by RNA-sequencing^16^. The copy number per cell values were averaged across three replicate amplified samples. The Pearson correlation coefficient is 0.91. All Pearson correlation coefficients were calculated with the log10 values. The dashed lines in (f) and (g) represent equality.

To determine the performance of MERFISH in amplified samples, we considered several performance metrics. First, we examined the average count per cell of the 10 barcodes not assigned to any RNAs, i.e. ‘blank’ barcodes. We found that 121 of the 130 RNA species in the 5×5 amplified MERFISH measurement had a higher copy number per cell than the maximum copy number per cell observed with the blank barcodes (Fig. 4d). A similar rate of ‘blank’ barcode detection was observed previously in unamplified samples^16^, indicating that amplification does not increase RNA misidentification rates. Next, we investigated the average 1-to-0 error and 0-to-1 error rate for each bit (Fig. 4e). We observed an average 1-to-0 error rate of ∼1.7% and a 0-to-1 error rate of ∼0.6%, comparable to the values observed previously with unamplified data^15^.

In addition, we found that the copy number per cell results were highly reproducible between replicates of MERFISH experiments with amplification (Fig. 4f). Next, to determine if amplification resulted in a decreased ability to detect RNAs, we compared the average copy number per cell from 5×5 amplified data with that from previously unamplified MERFISH data^15^, with the measured values per gene averaged across three replicates of amplified and unamplified samples. We found that these values correlated strongly with a Pearson correlation coefficient of 0.98 (Fig. 4g; ρ10 = 0.98 for the 123 RNA species whose measured copy numbers were larger than that observed for the largest blank barcode count) and that the average ratio was 1.04±0.03 (SEM, n = 123 RNAs). Thus, we conclude that amplification maintained the high detection efficiency (∼95%) previously reported for MERFISH^15^.

In addition, we compared the average copy number per cell detected for these RNAs by 5×5 amplified MERFISH measurements with the RNA abundance measured by RNA-seq (Fig. 4h). We observed a Pearson correlation coefficient of 0.91, comparable to our previously published data from unamplified samples^16^. Thus, based on each of these performance metrics, we conclude that bDNA amplification substantially increases the brightness of individual molecules measured with MERFISH without introducing additional noise.

### Amplification improves the performance of MERFISH for measurements with fewer encoding probes

The brightness of RNA signals in unamplified MERFISH measurements is set by the number of unique encoding probes targeted to each RNA; thus, amplification of MERFISH signals should allow the number of encoding probes to be reduced per RNA, which in turn would allow shorter genes to be targeted with MERFISH. However, it is worth noting that decreasing the number of encoding probes per gene will both decrease the average brightness of individual RNAs and increase the probability that a given RNA will, stochastically, not bind any encoding probes. Both of these effects are expected to decrease the efficiency of RNA detection with MERFISH; however, signal amplification is expected to only overcome the challenges introduced with the decrease in average signal brightness. Thus, we anticipate that the use of amplification will increase the detection efficiency in these cases but may not increase it to 100%.

To test the ability of bDNA amplification to improve the performance of MERFISH in circumstances where fewer encoding probes are used per gene, we designed an encoding probe library targeting the same 130 genes utilized above but in which we included only 16 encoding probes for each gene as opposed to the 92 utilized above. We have previously shown that the detection efficiency of MERFISH experiments is ∼100% when 92 encoding probes are used per gene. This experimental design thus allowed us to compare the results with the 92-probe experiments to quantify detection efficiency. We first performed MERFISH measurements with encoding 16 probes per gene in U-2 OS cells (Fig. 5a, b) without bDNA amplification. As expected, we observed that the signals from individual RNA molecules in each round of staining and imaging were indeed substantially dimmer than the signals observed when 92 encoding probes were used per gene. Nonetheless, we found that the average copy number per cell for individual RNAs correlated well with that measured using 92 encoding probes per gene (Fig. 5c), with a Pearson correlation coefficient of 0.8. However, we detected fewer RNA molecules per cell in these measurements: the ratio of the total RNA copy number per cell measured with 16 encoding probes relative to 92 encoding probes was 0.32. Across three independent replicates, this ratio was 0.33±0.02 (SEM) (Fig. 5g), confirming that the dimmer signals from individual RNA molecules led to fewer RNAs being detected.

**Figure 5.**
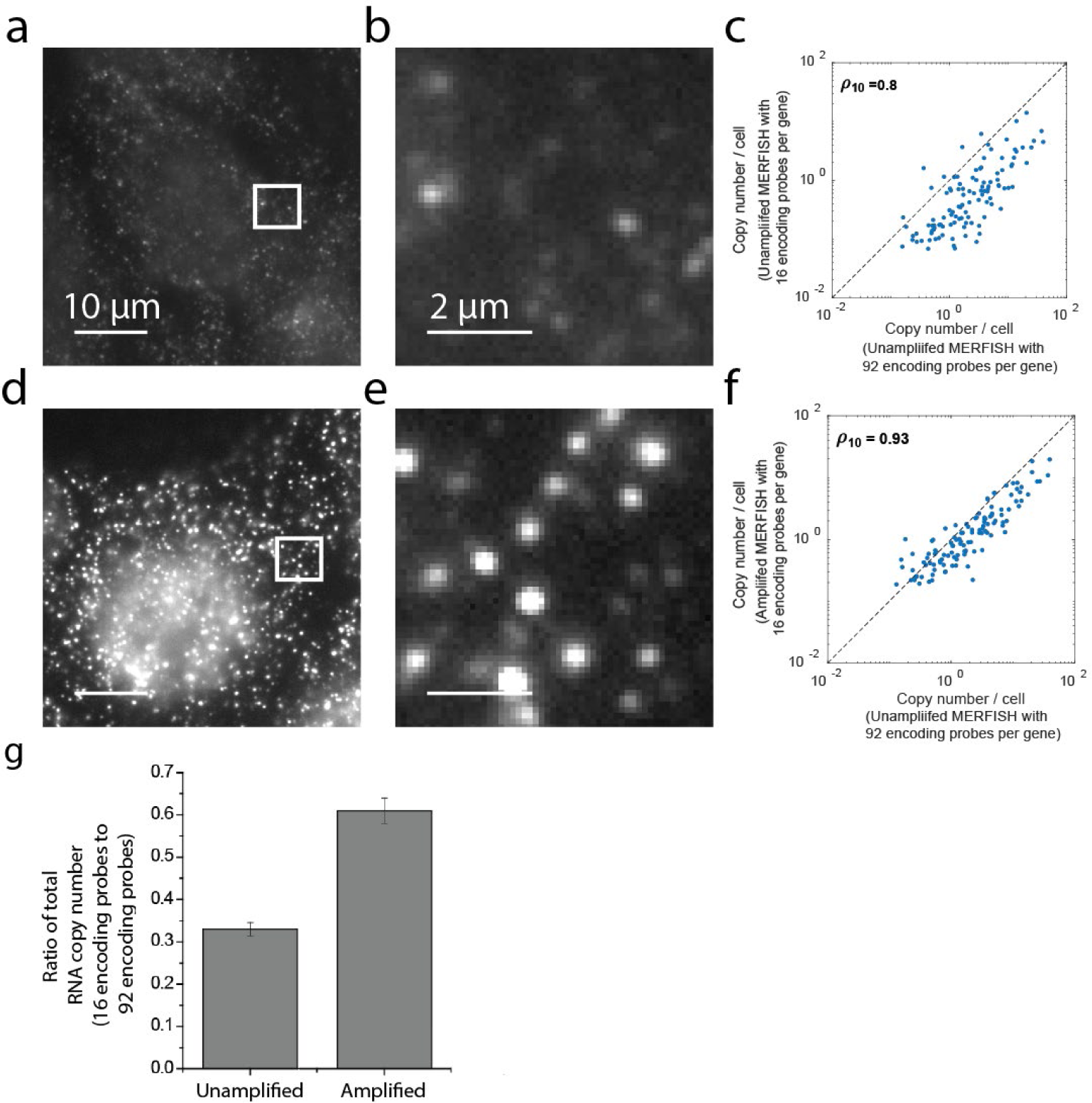
MERFISH measurements of 130 RNAs using only 16 encoding probes per gene. (a) Image of a U-2 OS sample stained with 16 encoding probes per gene and then stained with a single readout probe labeled with Cy5. Scale bar, 10 µm. No bDNA amplification was applied to this sample. (b) Zoom in of the white boxed region in (a). Scale bar, 2 µm. (c) The average copy number per cell observed for each of these RNA species as measured with 16 encoding probes per gene versus that measured with 92 per encoding probes per gene. Neither measurement was performed with bDNA amplification. The Pearson correlation coefficient between the two measurements is 0.8. The dashed line represents equality. (d, e) As in (a, b) but for a sample stained with 16 encoding probes per gene and amplified using the 5×5 bDNA approach. The contrast is the same as in (a, b). (f) As in (c) but with copy numbers per gene determined from 16 encoding probes per gene with bDNA amplification. The Pearson correlation coefficient is 0.93. (g) The ratio of the total RNA copy number measured with the amplified or unamplified 16-encoding-probe MERFISH measurement relative to the unamplified 92-encoding probe measurement. Error bars represent standard deviation across three replicate measurements.

To then determine the improvement provided with bDNA amplification, we repeat the MERFISH measurements with 16 encoding probes per gene in combination with our 5×5 bDNA amplification approach (Fig. 5d, e). As expected, amplification increased the brightness of these signals relative to the unamplified measurements. This increased brightness produced an increase in the correlation between the average copy number per cell determined with amplification to that determined with 92 encoding probes per gene but no amplification (Fig. 5f). In addition, the ratio of the total RNA copy number measured with the amplified 16-encoding-probe measurement relative to the unamplified 92-encoding-probe measurement was increased substantially. Across three replicates, this ratio was measured to be 0.61±0.03 (SEM) (Fig. 5g), indicating that amplification improved the detection efficiency of the 16-encoding-probe measurements by about 2-fold.

Because a minimum RNA length is required to accommodate a given number of encoding probes per gene, the use of fewer numbers of encoding probes per gene will allow MERFISH to target shorter genes, and bDNA amplification is likely to be highly beneficial for these measurements. Moreover, we anticipate that with improved encoding probe design, hybridization conditions, and other advances, it should be possible to increase the efficiency of encoding probe binding and further increase the detection efficiency of MERFISH for shorter genes.

### Discussion

We have presented here a combination of MERFISH and bDNA amplification. We showed that the bDNA amplifiers composed of three of the four nucleotides bind to targets rapidly, reaching saturated binding within 15 minutes. We demonstrated that this approach amplified a RNA smFISH signal by 5.5-fold, 10.5-fold, and 30.6-fold using 4×4 amplification, 5×5 amplification and 9×9 amplification, respectively, without increasing the size of fluorescent spots or increasing the variation in brightness from molecule to molecule, properties that are important for MERFISH performance. We also demonstrated that with bDNA amplification MERFISH maintained the ability to accurately identify and count RNAs with a near 100% detection efficiency with 92 encoding probes per gene. Finally, we showed that bDNA amplification substantially improved the detection efficiency of MERFISH (from ∼30% to ∼60%) when only 16 encoding probes were used per gene.

Notably, we observed a maximum amplification around 40% of the theoretical maximum for all amplifier designs (4×4, 5×5, and 9×9) tested. Thus, we suspect that the binding efficiencies of the amplifiers were not 100%. If we assume that the binding efficiencies were equal for each round of amplifier staining and readout staining, the average binding efficiency per round would have been ∼75%. In addition, we found that of the screened amplifier sequences, 20% of the amplifiers had either a high RNA-dependent background or a low amplification efficiency. In particular, the RNA-dependent background was unexpected because the sequences were blasted to avoid homology with the human transcriptome. We found that these specific amplifiers contained guanine nucleotides in the 20-mer repeating sequences that might form hybrid G-quadruplex structures, which have been reported to bind to mitochondrial RNA^33^. The specific spatial distribution observed with the staining of these amplifiers (Fig. 3a, b) was consistent with the distribution of mitochondria, supporting this hypothesis. In parallel, the low amplification efficiency of a few amplifiers could have arisen for a few reasons. One reason could be that the melting temperatures (Tm) of amplifier sequences were unexpectedly low, inhibiting the assembly of these amplifiers or promoting their disassembly in the hybridization and wash conditions used. Alternatively, it is possible that these amplifiers formed stable G-quadruplex structures, which inhibited the binding of those secondary amplifiers to primary amplifiers. Despite these few unexpected failures, the vast majority of the designed amplifiers worked as anticipated, suggesting that this bDNA approach could be extended to the amplification of more readout sequences and hence to MERFISH measurement with longer barcodes, detecting more RNAs. We also anticipate that it is possible to design amplifiers to allow a third or fourth round of amplification and produce signal intensities ∼100-1000-fold larger.

The substantial increase in signal brightness generated by bDNA amplification should greatly facilitate several areas of applications of MERFISH. First, we have performed MERFISH measurements using 16 encoding probes per RNA. Because only a fraction of the designed encoding probes actually bind to each RNA, encoding probes can be designed such that they target overlapping regions of the same RNA without reducing the number of probes actually bound to each RNA molecule^14, 34^. Thus, with this probe design, we anticipate that RNAs as short as a few hundred nucleotides can be detected with MERFISH. With additional improvements in encoding probe design and hybridization conditions, it may be possible to detect even shorter genes. Moreover, the ability to detect RNA molecules with relatively few probes would also improve RNA isoforms discrimination. Second, with signal amplification, it should be possible to substantially reduce the imaging duration by using shorter exposure times. As the imaging time is typically a sizeable portion of the total time required to perform a MERFISH measurement, it should be possible to substantially improve MERFISH imaging throughput with amplification. Third, bDNA amplification could increase MERFISH performance for high background samples. For example, amplification will increase the signal above background sources such as autofluorescence in tissue samples that can be challenging to remove even with clearing approaches (e.g. lipofuscin^17^). Thus, bDNA amplification should greatly facilitate imaging of tissues with high autofluorescence background. Finally, amplification could substantially reduce the cost of MERFISH measurements, including lowering the requirement of high-power lasers on microscopy setups and reducing the number of probes required per gene in a MERFISH library. Thus, we anticipate that the improved MERFISH performance and versatility provided by this amplification method should facilitate the application of spatially resolved single-cell transcriptomics to a wide array of biological questions.

## Methods

### Design of the encoding probes

MERFISH measurements in human osteosarcoma cells (U-2 OS) (ATCC) were performed with the same MERFISH-encoding probe set as previously described^16^. Briefly, a 16-bit MHD4 code was used to encode the RNAs. In this encoding scheme, each of the 140 possible barcodes has a constant Hamming weight (i.e., the number of “1” bits in each barcode) of 4 to avoid potential bias in the measurement of different barcodes due to a differential rate of “1” to “0” and “0” to “1” errors. In addition, all barcodes have a Hamming distance of at least 4 to enable error detection and error correction. We used 130 of the 140 possible barcodes to encode cellular RNAs and the remaining 10 barcodes were not assigned to an RNA and served as blank controls. In the first encoding probe library, each RNA species had 92 encoding probes, with each encoding probe containing three of the four readout sequences assigned to each RNA. Our second encoding probe library was designed identically to the first with the distinction that each RNA species had 16 encoding probes. The encoding probe set targeting FLNA (Biosearch) was described previously^16^.

Encoding probe construction, coverslip silanization, cell culture and fixation, encoding probe staining, and gel embedding and clearing were performed as previously described^15^.

### Amplifier staining

Primary and secondary amplifiers were stained in gel embedded and cleared samples. Samples were incubated for 5 min in a 10% formamide wash buffer, containing 2× SSC (ThermoFisher) and 10% (vol/vol) formamide (ThermoFisher) in nuclease-free water. Next, 50 μL of 5 nM each of the primary amplifiers (Integrated DNA Technologies) in amplifier hybridization buffer, containing 2× SSC, 10% (vol/vol) formamide, 0.1% (wt/vol) yeast tRNA (Life Technologies), 1% (vol/vol) murine RNase inhibitor (New England Biolabs), and 10% (wt/vol) dextran sulfate (Sigma), was added to a Parafilm-coated surface to form droplets. Samples were inverted onto these 50 μL droplets after removing extra 10% formamide wash buffer with Kimwipes and incubated in a humidity-controlled 37 °C incubator for 30 minutes (for all MERFISH measurements) or for 15 minutes, 30 minutes, 60 minutes and 180 minutes for the binding rate measurements. Then the samples were washed three times in 10% formamide wash buffer at room temperature in Petri dishes for 5 minutes each. To bind the secondary amplifiers, 50 μL of 5 nM each of the secondary amplifiers (Integrated DNA Technologies) in the same amplifier hybridization buffer described above was placed on a fresh parafilm-coated surface, the samples were inverted onto these droplets, and the hybridization was performed as described for the primary amplifiers. Then the samples were washed twice in 10% formamide wash buffer at room temperature for 5 minutes each followed by a third wash in 10% formamide wash buffer in 37 °C incubator for 15 minutes.

The samples were either imaged immediately or stored in 2× SSC supplemented with 0.1% (vol/vol) murine RNase inhibitor at 4 °C for no longer than 48 h. Readout probes were hybridized to these samples as described previously^15^.

### MERFISH imaging platforms

The samples were imaged on a home-built high-throughput imaging platform at the Center for Advanced Imaging, Harvard University. Briefly, this microscope was constructed around a Nikon Ti Eclipse microscope body and a Nikon, CFI Plan Apo Lambda 60x oil objective. Illuminations in 750, 647, 560, 488 and 405 nm were provided using solid-state lasers (MBP Communications, 2RU-VFL-P-500-750-B1R; MBP Communications, 2RU-VFL-P-2000-647-B1R; MBP Communications, 2RU-VFL-P-2000-560-B1R; MBP Communications, 2RU-VFL-P-500-488-B1R; Coherent, Cube 405). These laser lines were used to excite readout probes labeled with Alexa750 and Cy5, orange fiducial beads, a Poly dT readout probe (Alexa 488) and DAPI, respectively. The illumination profile was flattened with a πShaper (Pishaper). The fluorescence emission from the sample was separated from the laser illumination using a penta-band dichroic (Chroma, zy405/488/561/647/752RP-UF1) and imaged with a scientific CMOS camera (sCMOS; Hamamatsu, C11440-22CU) after passing through two duplicate custom penta-notch filters (Chroma, ZET405/488/561/647–656/752m) to remove stray excitation light. The pixel size for the sCMOS camera corresponded to 109 nm in the sample plane. The exposure time was 500 ms for each imaging frame. Sample X/Y position was controlled via a motorized microscope stage (Ludl). Sample focus was maintained by feedback on the reflection of an IR laser (Thorlabs, LP980-SF15) off the sample coverslip interface. The reflected IR signal was detected by a CMOS camera (Thorlabs, DCC1545M) and the sample to objective distance was controlled by an objective nanopositioner (Mad City Labs, Nano-F100S).

The sample coverslip was held inside a flow chamber (Bioptechs, FCS2), and buffer exchange within this chamber was directed using a custom-built automated fluidics system described previously^9^, controlling three eight-way valves (Hamilton, MVP and HVXM 8–5) and a peristaltic pump (Gilison, Minipuls 3).

### Sample imaging

Sequential MERFISH imaging was carried out on the high-throughput imaging platform descripted in the *MERFISH imaging platforms* section. Briefly, each MERFISH round consisted of readout probes staining (10 min), wash buffer washing (5 min), imaging buffer exchange (3 min), imaging of 100-400 different fields of view (FOV), readout fluorophore cleavage (15 min), and 2xSSC washing (5 min). 8 rounds of two color MERFISH imaging were performed on each sample. Specifically, the readout probes staining was done by flowing 3 nM readout probes in hybridization buffer comprised of 2× SSC, 10% (vol/vol) ethylene carbonate (Sigma-Aldrich), 0.1% (vol/vol) murine RNase inhibitor (NEB) in nuclease-free water. Wash buffer contained 2× SSC and 10% (vol/vol) ethylene carbonate in nuclease-free water. Imaging buffer was made of 2× SSC, 50 mM Tris·HCl pH 8, 10% (wt/vol) glucose, 2 mM Trolox (Sigma-Aldrich), 0.5 mg/mL glucose oxidase (Sigma-Aldrich), 40 μg/mL catalase (Sigma-Aldrich), and 0.1% (vol/vol) murine RNase inhibitor in nuclease-free water. Cleavage buffer comprising 2× SSC and 50 mM of Tris (2-carboxyethyl) phosphine (TCEP; Sigma) was used to cleave the disulfide bond conjugating dyes to the probes. smFISH imaging was performed using these protocols but without the cleavage or 2xSSC washing steps. Each MERFISH measurement contained ∼1000 cells.

### Image processing and decoding

For FLNA smFISH data, the brightness and PSF sizes of unamplified and amplified FISH spots were calculated via a Gaussian fitting routine previously described^35^.

For the MERFISH data, registration of images of the same FOV across different imaging rounds as well as decoding of the RNA barcodes was conducted using a previous analysis pipeline^16^. Briefly, the drift between images in each imaging round was corrected using the localizations of the fiducial beads in each round of imaging. Next, background in the images were removed via a high-pass filter, and RNA spots were tightened by deconvolution. We have previously found that the signal from the same RNA can vary slightly in position (<100 nm) from round to round. To address this small variation, we then low-pass filtered each image to ensure the signal from the same RNA overlapped in different imaging rounds. To remove the natural variation in brightness between color channels and different imaging rounds, we normalized the intensity measured for each color channel in each imaging round via the 95% quantile of the corresponding brightness histogram, effectively equalizing the brightness histograms observed for different bits. We then compared the normalized intensity for a given pixel in each of the 16 images to the expected intensity produced by each barcode for each of the 16 bits, and we selected the barcode that best matched the observed intensity pattern for that pixel. However, if the Euclidean distance between this pixel intensity pattern and the expected intensity pattern from the closest barcode was larger than a maximum threshold value, the pixel was not assigned to a barcode. This threshold distance was set by the maximum Euclidean distance between a correct barcode and each of the incorrect barcodes generated by flipping a single bit in that correct barcode. Finally, adjacent pixels assigned to the same barcode were then combined to form a putative RNA.

This pipeline was run on a desktop server that contained two 10-core Intel Xeon E5-2680 2.8-GHz CPUs and 256 GB of RAM.

## Data availability

The datasets that support the findings of this paper are available from the corresponding authors upon request.

## Code availability

The software used to analyze the datasets are available from the corresponding authors upon request.

## Competing interests

The authors are inventors on patents applied for by Harvard University that describe MERFISH related technologies.

## Author Contributions

C.X., J.R.M., and X.Z designed the experiments, C.X. conducted experiments and analyzed data, C.X. and H.P.B. contributed reagents and custom instrumentation, C.X., J.R.M., and X.Z. wrote the manuscript with input from H.P.B. All authors reviewed the manuscript.

